# Inactivation of the core *cheVAWY* chemotaxis genes disrupts chemotactic motility and organised biofilm formation in *Campylobacter jejuni*

**DOI:** 10.1101/449850

**Authors:** Mark Reuter, Eveline Ultee, Yasmin Toseafa, Andrew Tan, Arnoud H.M. van Vliet

**Affiliations:** Gut Health and Food Safety Programme, Quadram Institute Bioscience, Norwich Research Park, Norwich, United Kingdom; Department of Pathology and Infectious Diseases, School of Veterinary Medicine, Faculty of Health and Medical Sciences, University of Surrey, Guildford, United Kingdom

**Keywords:** Campylobacter, chemotaxis, biofilm formation, flagellar motility

## Abstract

Flagellar motility plays a central role in the bacterial foodborne pathogen *Campylobacter jejuni*, as flagellar motility is required for reaching the intestinal epithelium and subsequent colonisation or disease. Flagellar proteins also contribute strongly to biofilm formation during transmission. Chemotaxis is the process directing flagellar motility in response to attractant and repellent stimuli, but its role in biofilm formation of *C. jejuni* is not well understood. Here we show that inactivation of the core chemotaxis genes *cheVAWY* in *C. jejuni* strain NCTC 11168 affects both chemotactic motility and biofilm formation. Inactivation of any of the core chemotaxis genes (*cheA*, *cheY*, *cheV* or *cheW*) impaired chemotactic motility but did not affect flagellar assembly or growth. The Δ*cheY* mutant swam in clockwise loops, while complementation restored normal motility. Inactivation of the core chemotaxis genes interfered with the ability to form a discrete biofilm at the air-media interface, and the Δ*cheY* mutant displayed reduced dispersal/shedding of bacteria into the planktonic fraction. This suggests that while the chemotaxis system is not required for biofilm formation *per se*, it is necessary for organized biofilm formation. Hence interference with the *Campylobacter* chemotaxis system at any level disrupts optimal chemotactic motility and transmission modes such as biofilm formation.

## INTRODUCTION

*Campylobacter jejuni* is an important causative agent of bacterial gastroenteritis in humans (Nichols *et al*, 2012), and is commonly transmitted via contaminated food, especially poultry meat (Tam *et al*, 2009). Infection with *C. jejuni* is also associated with neurodegenerative diseases like Miller-Fisher and Guillain-Barré syndrome (Poropatich *et al*, 2010). One of the key factors for *C. jejuni* is its flagella-based motility, as aflagellated *C. jejuni* are unable to cause disease in animal models, show strongly reduced host cell invasion in cell culture-based assays (Gao *et al*, 2014; Lertsethtakarn *et al*, 2011), are targeted by bacteriophages (Baldvinsson *et al*, 2014). and show reduced biofilm formation (Brown *et al*, 2014; Reuter *et al*, 2010; Svensson *et al*, 2014).

Biofilms are bacterial populations encased in an extracellular matrix, and the biofilm environment can support survival of the bacterial population in adverse environmental conditions, and assist in resistance to antimicrobials such as disinfectants and antibiotics (Costerton *et al*, 1999; Teh *et al*, 2014). *C. jejuni* can form a monospecies biofilm but can also integrate into biofilms already present (Hanning *et al*, 2008; Reeser *et al*, 2007; Teh *et al*, 2019; Teh *et al*, 2010). *C. jejuni* mutants with defects in biofilm formation show reduced chicken colonization, adhesion and invasion of epithelial cells, as well as reduced intracellular survival (Rahman *et al*, 2014; Svensson *et al*, 2009; Theoret *et al*, 2012). While flagella are not required for biofilm formation in *C. jejuni*, flagellated strains show higher levels of biofilm formation, and *C. jejuni* cells isolated from biofilms show higher levels of expression of flagellar proteins (Kalmokoff *et al*, 2006; Reuter *et al*, 2010; Svensson *et al*, 2014).

In bacteria, motility is commonly directed in response to attractant and repellant external and intracellular signals, with the core chemotaxis proteins CheA and CheY transmitting the signal to the flagellar switch to alter flagellar motor rotation (Korolik, 2018; Lertsethtakarn *et al*, 2011). The signals are sensed by methyl-accepting chemotaxis proteins (MCPs), and this sensing initiates a signalling cascade that results in autophosphorylation of the histidine kinase CheA. The phosphorylated CheA transfers the phosphate group to the response regulator CheY, with phosphorylated CheY interacting with the flagellar switch to alter flagellar motor rotation. *C. jejuni* contains a number of accessory proteins, such as the CheW and CheV proteins, a methyltransferase (CheR) and a methylesterase (CheB) (Korolik, 2018; Lertsethtakarn *et al*, 2011), as well as a large number of MCPs involved in sensing different stimuli relevant to transmission and infection, such as amino acids, deoxycholate, dicarboxylic acid TCA intermediates, fucose, and nutrients (Day *et al*, 2016; Dwivedi *et al*, 2016; Lubke *et al*, 2018; Reuter and van Vliet, 2013). Chemotaxis-defective mutants have been shown to be attenuated in disease models (Chandrashekhar *et al*, 2015; Hendrixson and DiRita, 2004; Yao *et al*, 1997), show reduced immunopathology (Bereswill *et al*, 2011) and chick colonization (Hendrixson and DiRita, 2004).

It was previously shown that fucose utilisation and chemotaxis are linked to biofilm formation (Dwivedi *et al*, 2016), and biofilm formation in *C. jejuni* is responsive to external stimuli such as oxygen availability (Reuter *et al*, 2010; Teh *et al*, 2017). In a recent study, biofilm formation as well as autoagglutination was shown to be increased in *C. jejuni cheV* and *cheW* mutants (Tram *et al*, 2020), and although these mutants were not complemented, it suggests a more direct link between chemotaxis and biofilm formation in *Campylobacter*. Here we have investigated the role of additional chemotaxis components in *C. jejuni* biofilm formation and chemotactic motility, by inactivation and complementation of the individual core chemotaxis genes *cheVAWY* in *C. jejuni*, and qualitative assessment of biofilm formation at air-liquid interfaces.

## MATERIALS AND METHODS

### *C. jejuni* strains and growth conditions

*Campylobacter jejuni* strain NCTC 11168 and its isogenic mutants (Table 1) were routinely cultured in a MACS-MG-1000 controlled atmosphere cabinet (Don Whitley Scientific) in microaerobic conditions (85% N2, 10% CO_2_, 5% O_2_,) at 37°C. For growth on plates, strains were either grown on Brucella agar or Blood Agar Base (BAB) with Skirrow supplement (10 μg ml^−1^ vancomycin, 5 μg ml^−1^ trimethoprim, 2.5 IU polymyxin-B). Broth culture was carried out in Brucella broth (Becton Dickinson).

**Table 1.** *Campylobacter jejuni* strains used in this study.

| Strain | Description <sup>a</sup> |
| --- | --- |
| ***C. jejuni* strains** |  |
| NCTC 11168 | Wild-type *C. jejuni* (Parkhill *et al*, 2000) |
| 11168 $\Delta cheY$ | NCTC 11168 *cj1118c::kan<sup>R</sup>* |
| 11168 $\Delta cheW$ | NCTC 11168 *cj0283c::kan<sup>R</sup>* |
| 11168 $\Delta cheA$ | NCTC 11168 *cj0284c::cat<sup>R</sup>* |
| 11168 $\Delta cheV$ | NCTC 11168 *cj0285c::kan<sup>R</sup>* |
| 11168 $\Delta cheY + cheY$ | NCTC 11168 *cj1118c::kan<sup>R</sup> cj0046::cj1118c<sup>fdxApr\*</sup>cat<sup>R</sup>* |
| 11168 $\Delta cheW + cheW$ | NCTC 11168 *cj0283c::kan<sup>R</sup> cj0046::cj0283c<sup>fdxApr\*</sup>cat<sup>R</sup>* |
| 11168 $\Delta cheA + cheA$ | NCTC 11168 *cj0284c::cat<sup>R</sup> cj0046::cj0284c<sup>fdxApr\*</sup>kan<sup>R</sup>* |
| 11168 $\Delta cheV + che^*$ | NCTC 11168 *cj0285c::kan<sup>R</sup> cj0046::cj0285c<sup>fdxApr\*</sup>cat<sup>R</sup>* |
| 11168 $\Delta pflA$ | NCTC 11168 *cj1565c::kan<sup>R</sup>* (Reuter *et al*, 2015) |

a. *kan<sup>R</sup>*, kanamycin resistance cassette; *cat<sup>R</sup>*, chloramphenicol resistance cassette; \* denotes complementation construct: <sup>fdxApr</sup>, gene under control of the constitutive *fdxA* promoter.

### Insertional inactivation and complementation of chemotaxis genes

Insertional inactivation mutants were made as described previously (Reuter and van Vliet, 2013) using primers listed in Table S1, resulting in plasmids listed in Table S2. Plasmids were propagated in *E. coli* strain TOP10. To insert antibiotic resistance cassettes, *Bam*HI sites were introduced in the target genes by inverse PCR (Table S1) and ligated to either the kanamycin-resistance cassette (Δ*cheY*, Δ*cheW*, Δ*cheV* mutants) or chloramphenicol-resistance cassette (Δ*cheA* mutant). All constructs were sequenced prior to transformation (Eurofins Genomics, Ebersberg, Germany). Single *C. jejuni* mutant strains (Table 1) were isolated after transformation of the *C. jejuni* NCTC 11168 wild-type strain with plasmids by electroporation (Reuter and van Vliet, 2013), followed by selection on plates supplemented with either 50 μg ml^−1^ kanamycin or 10 μg ml^−1^ chloramphenicol. To confirm the position of the antibiotic resistance cassette in antibiotic resistant clones, genomic DNA was isolated from four ml of overnight culture (DNeasy kit, QIAGEN).

Diluted genomic DNA (50 ng) was used as template for PCR using primers that anneal outside of the cloned flanking regions in combination with antibiotic resistance cassette-specific primers (Table S1). *C. jejuni* mutants were complemented (Table 1) by inserting the individual chemotaxis genes (*cheA*, *cheY*, *cheW*, *cheV*) *in trans* using the *cj0046* pseudogene, as described previously (Reuter and van Vliet, 2013). In these plasmids, the genes are expressed from the *fdxA* promoter (van Vliet *et al*, 2001).

### Light microscopy and flagella staining using the Ryu stain

Typically, 10 μl of an overnight culture was examined using an Eclipse 50i microscope (×100 lens) to monitor swimming phenotypes. When necessary, a Coolpix 4500 digital camera (Nikon) was used to capture video (15 frames second^−1^, 320×240 pixels). Video compilations were made by extracting appropriate frames using ImageJ (Rasband, W.S., ImageJ, U. S. National Institutes of Health, Bethesda, Maryland, USA, http://imagej.nih.gov/ij/, 1997-2014). To plot swimming behaviour, individual cells were tracked using the Manual tracking plug-in for ImageJ (Fabrice Cordeli, http://rsb.info.nih.gov/ij/plugins/track/track.html) then visualized using the Chemotaxis and Migration Tool (Ibidi, http://ibidi.com/software/chemotaxis_and_migration_tool/). To visualise flagella, cells were stained using the Ryu stain (Heimbrook *et al*, 1989) as described previously (Reuter *et al*, 2015). ImageJ was used to add a scale bar and prepare montage images.

### Chemotaxis assays

Chemotactic motility was measured using soft agar motility assays and tube taxis assays (Reuter and van Vliet, 2013). All soft agar motility assays were carried out using Brucella soft agar in square 10 mm^2^ petri plates (Sterilin) inoculated with wild-type and three test strains, as described previously (Reuter and van Vliet, 2013). For each plate, halo size was expressed as a percentage of the corresponding wild-type and each strain was tested for significance using a one-sample t-test (alpha = 0.05), compared to a hypothetical value of 100 (GraphPad Prism 6.01). Strain-to-strain comparisons were made using a two-tailed Mann-Whitney test.

Tube taxis assays were prepared as described previously using Brucella soft agar (Reuter and van Vliet, 2013). The tubes were incubated at 37°C in air in a waterbath (Grant). Tubes were photographed after 24, 40, 48, 64, and 72 hours and the dye front was measured from the top of the agar using the ImageJ software and expressed as a percentage of the wild-type strain. Each strain was tested for significance using a one-sample t-test (alpha = 0.05) compared to a hypothetical value of 100 (GraphPad Prism 6.01) and strain-to-strain comparisons were made using a two-tailed Mann-Whitney test.

### Biofilm assays

A 50 μl single-use glycerol stock, routinely stored at −80°C, was used to inoculate a BAB plate with Skirrow supplements and these cells were used to inoculate fresh Brucella broth. Cultures were grown in microaerobic conditions with shaking overnight at 37°C. The overnight culture was diluted to A_600_ ≈ 0.05 (~ 1 x 10^7^ CFU ml^−1^) in 22 ml sterile Brucella in a 50 ml falcon tube. A twin-frost microscope slide (sterilized in 70% ethanol) was inserted into each falcon tube. Tubes were incubated at 37°C in microaerobic conditions without shaking. After 48 hours, slides were removed from the tubes using flamed-sterilized tweezers and briefly washed in water. Slides were dried in air before staining with 1% crystal violet. Unbound crystal violet was washed off with water and slides were dried in air. Biofilms were imaged using a GenePixPro microarray scanner (Axon). The photomultiplier tube (PMT) gain of either the 635 nm or 532 nm laser was adjusted to achieve a balanced image. To assess biofilm shedding, the A_600_ of the planktonic cultures were measured; statistically different results were determined using a two-tailed Mann-Whitney test (GraphPad Prism 6.01).

## RESULTS

### Inactivation and complementation of the genes involved in the core chemotaxis system

To assess the roles of the *cheVAWY* genes in *C. jejuni* chemotaxis, each individual gene was inactivated by a kanamycin or chloramphenical antibiotic resistance cassette in the same orientation as the gene (Table 1). To test for polar effects, each mutated gene was complemented by introduction of the different chemotaxis genes mutant *in trans* in the *cj0046* pseudogene under control of the *fdxA* promoter (Reuter and van Vliet, 2013). Inactivation of the chemotaxis genes did not result in significant changes in growth (Figure S1), and all isolates expressed flagella at both poles (Fig. 1). As the *cheV*, *cheA* and *cheW* genes are in an operon (Dugar *et al*, 2013; Porcelli *et al*, 2013), we confirmed that the downstream *cheAW* or *cheW* genes were still transcribed by reverse-transcriptase PCR in the Δ*cheV* and Δ*cheA* mutants, respectively (not shown), although we cannot estimate changes in expression levels using this experimental approach. For all mutants and complemented strains described here, chemotactic motility was assessed using soft agar motility plates, and using tube taxis assays which measure motility in both an energy and redox gradient (Reuter and van Vliet, 2013).

**Figure 1.**
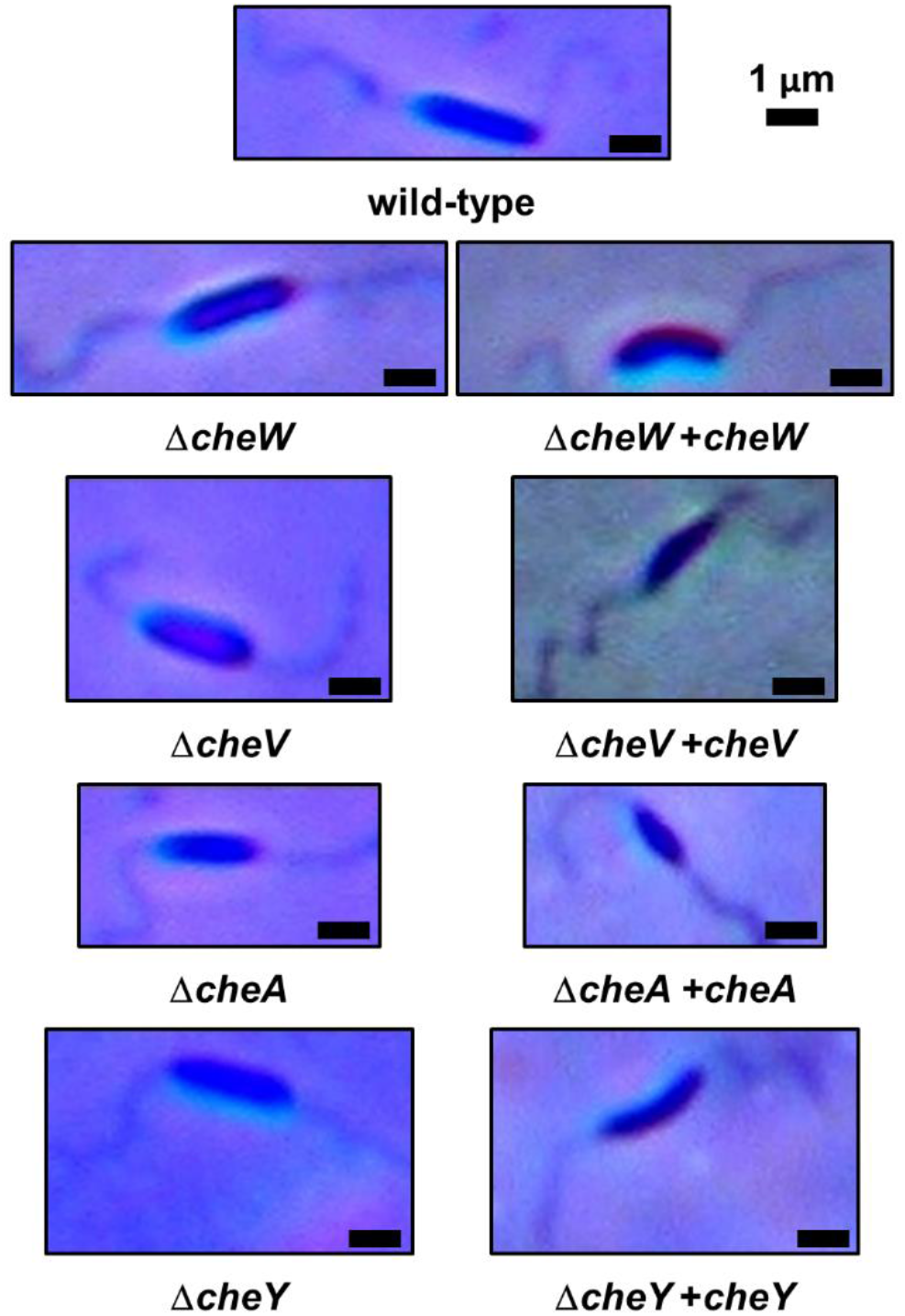
Disrupting chemotaxis proteins does not impair flagella assembly. Brucella broth cultures of wild-type *C. jejuni* NCTC11168 and the chemotaxis gene mutants and complemented strains were grown overnight at 37°C in microaerobic conditions. Cell were mounted between a microscope slide and coverslip and freshly prepared Ryu stain added adjacent to the coverslip. After five minutes, areas where the Ryu stain had penetrated were photographed at ×100 magnification using a Nikon Coolpix 4500 digital camera. Montage images and scale bars were prepared using ImageJ. Scale bar = 1 micron. Pictures are representative examples of multiple cultures each examined.

### CheV is the dominant adaptor protein in the chemotaxis signalling pathway

In liquid media, motility of the Δ*cheW* and Δ*cheV* mutants was comparable to the wild-type, but both halo formation and migration were reduced compared to the wild-type (Figures 2 and 3). Chemotactic motility of the Δ*cheW* mutant was reduced to less than 50% compared to the wild-type strain at both 24 and 48 hours, respectively. The Δ*cheV* mutant showed greater reduction in halo formation and migration in soft agar than the Δ*cheW* mutant, when compared to the wild-type strain at both 24 and 48 hours. Complementation of the Δ*cheW* mutant restored chemotaxis phenotypes to that of the wild-type strain. However, complementation of the Δ*cheV* mutant did not restore chemotactic motility.

**Figure 2.**
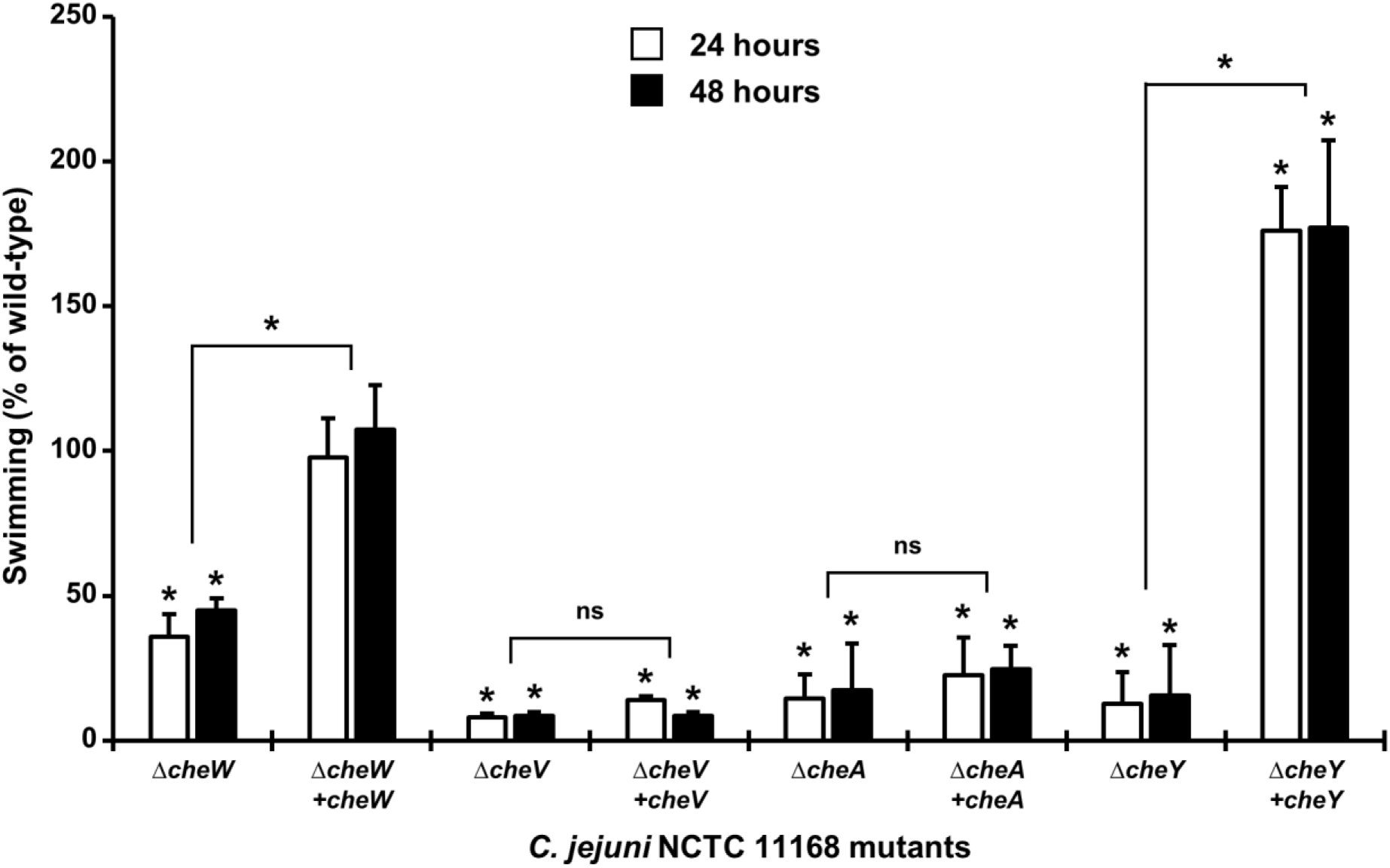
Chemotaxis mutants show a significant reduction in chemotactic motility in soft agar. For each chemotaxis mutant, the mutant strain was inoculated into 0.4% agar with the wild-type strain. Halo formation was measured after 24 and 48 hours and halo area expressed as the percentage of the halo for the wild-type strain. Raw mean area of WT was 293.2 ± 15.52 and 1230 ± 63.69 mm^2^ (24 and 48 hrs). Results are the mean from at least three biological replicates and error bars show standard deviation. An asterisk denotes statistically significant results, based on a one-sample *t*-test (comparison with wild-type) or a two-tailed Mann-Whitney test (p < 0.05). NS represents non-significant results.

**Figure 3.**
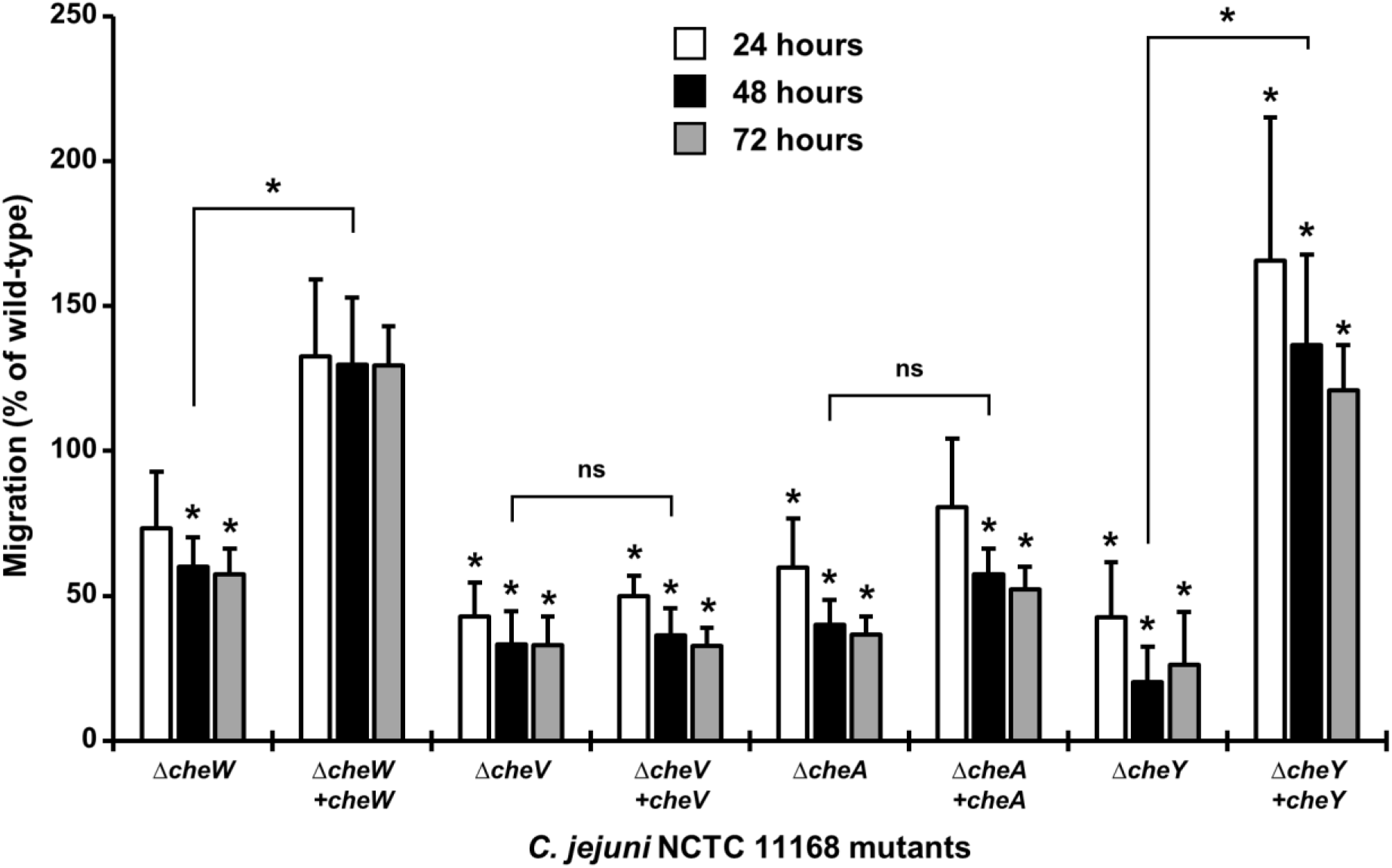
Chemotaxis mutants show a significant reduction in migration in an energy and redox gradient. Tube taxis assays for each strain were monitored over 72 hours at 37°C and the dye front measured after 24, 48 and 72 hours. Migration, calculated as a percentage of the wild-type used in the same assay, is shown at 24, 48, and 72 hrs compared to WT (100%) is shown. Raw mean migration for WT was 8.198 ± 0.981, 25.58 ± 2.94 and 48.03 ± 5.09 mm for 24h, 48h, and 72h, respectively. Results are the mean from at least four biological replicates and error bars show standard deviation. Asterisks denote statistically significant results, based on a one-sample *t*-test comparing mutant with wild-type (p < 0.05). NS represents non-significant results.

### Inactivation of the core chemotaxis genes*cheY* and*cheA* disrupts chemotactic motility

The Δ*cheY* mutant displayed the previously described chemotaxis defect (Yao *et al*, 1997), with chemotactic motility reduced to less than 20% of wild-type (Figures 2 and 3). Chemotactic motility was restored to wild-type levels by complementation with the *cheY* gene, and even exceeded that observed in the wild-type strain. The Δ*cheY* mutant showed a swimming behaviour reminiscent to the ‘catherine wheel’ firework (Lauga *et al*, 2006) in liquid media, where a cell would appear to get trapped in a clockwise swimming loop (see Supplementary Movie 1 and Fig. 4B) before resuming a darting motility as observed in the wild-type (Fig. 4A), while complementation of the Δ*cheY* mutant restored wild-type swimming behaviour (Fig. 4C). Inactivation of the *cheA* gene strongly reduced chemotactic motility, but introduction of the *cheA* gene *in trans* did not complement the Δ*cheA* mutant (Figures 2 and 3), suggesting that other factors such as expression levels and stoichometry may be important in CheA function.

**Figure 4.**
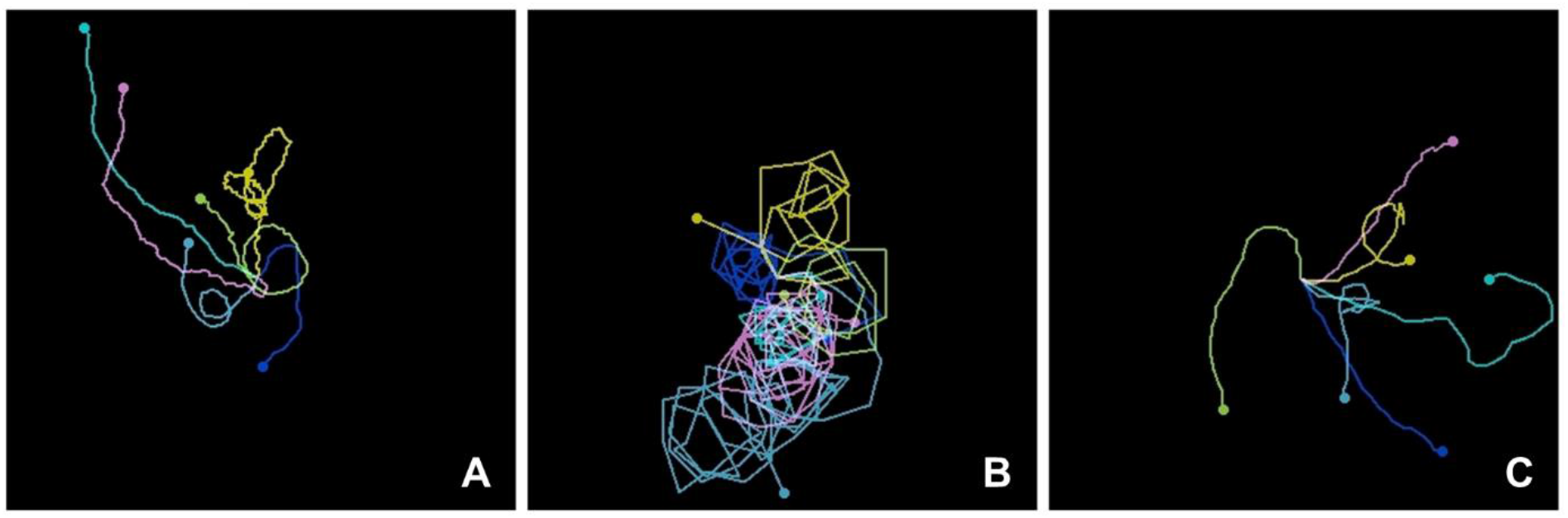
The Δ*cheY* mutant shows a swimming phenotype consisting of repeated clockwise swimming, similar to a ‘catherine wheel’ fireworks. Following 6 – 7 hours growth at 37°C in microaerobic conditions, movies of swimming cells (A, wild-type; B, Δ*cheY*; C, Δ*cheY +cheY*;) were recorded at ×100 magnification using a light microscope and attached Nikon camera. Individual cells were tracked using the Manual Tracking ImageJ plug-in and swimming trajectories plotted using the Chemotaxis and Migration Tool (Ibidi).

### Chemotaxis is required for organized biofilm formation at the air-media interface

We investigated the role of chemotaxis in biofilm formation, by developing a glass slide-based assay combined with crystal violet staining, with detection using a microarray scanner (employing both lasers at 532 and 635 nm). The wild-type strain formed a biofilm on the glass slide at the air-media interface (Fig. 5). Below the air-media interface, a less-intense ~2 mm band of adhered cells was observed. All of the chemotaxis mutants used in this study were defective in formation of a biofilm at the air-media interface (Fig. 5). In the Δ*cheW* mutant, the air-media interface biofilm was present, but a second more intense area of adhered cells was also visible ~5 mm below the air-media interface. Both the Δ*cheV* mutant and the complemented Δ*cheV* mutant showed defective air-media interface biofilm. The Δ*cheA* and Δ*cheY* mutants both showed a disorganized air-media interface biofilm with a greater population of submerged cells. In all cases, except for the *cheV* and in part the *cheA* mutant, complementation with the respective wild-type gene restored the organised interface biofilm phenotype (Fig. 5). To assess the level of shedding of cells from the biofilm, the A_600_ of planktonic fraction for each assay was recorded. Inactivation of *cheY* resulted in fewer cells in the planktonic fraction compared to the wild-type strain, while the complemented Δ*cheY* mutant had more cells in the planktonic fraction (Fig. S2). Therefore, the biofilm phenotype of at least then Δ*cheY* mutant could in part be due to a disruption of normal cell shedding/dispersal from the biofilm. To test the hypothesis that *C. jejuni* may require the fully functional chemotactic motility to seek the optimal environment for growth as a biofilm, the slide biofilm assay was performed using a paralysed flagella (Δ*pflA*) mutant, which expresses flagella but is non-motile (Reuter *et al*, 2015; Yao *et al*, 1994). This mutant displayed low levels of biofilm formation at the air-media interface (Fig. 5). Thus, while the chemotaxis system is not required for biofilm formation *per se*, it is necessary for organized biofilm formation at the air-media interface and may play a role in dispersal/shedding of cells from the biofilm.

**Figure 5.**
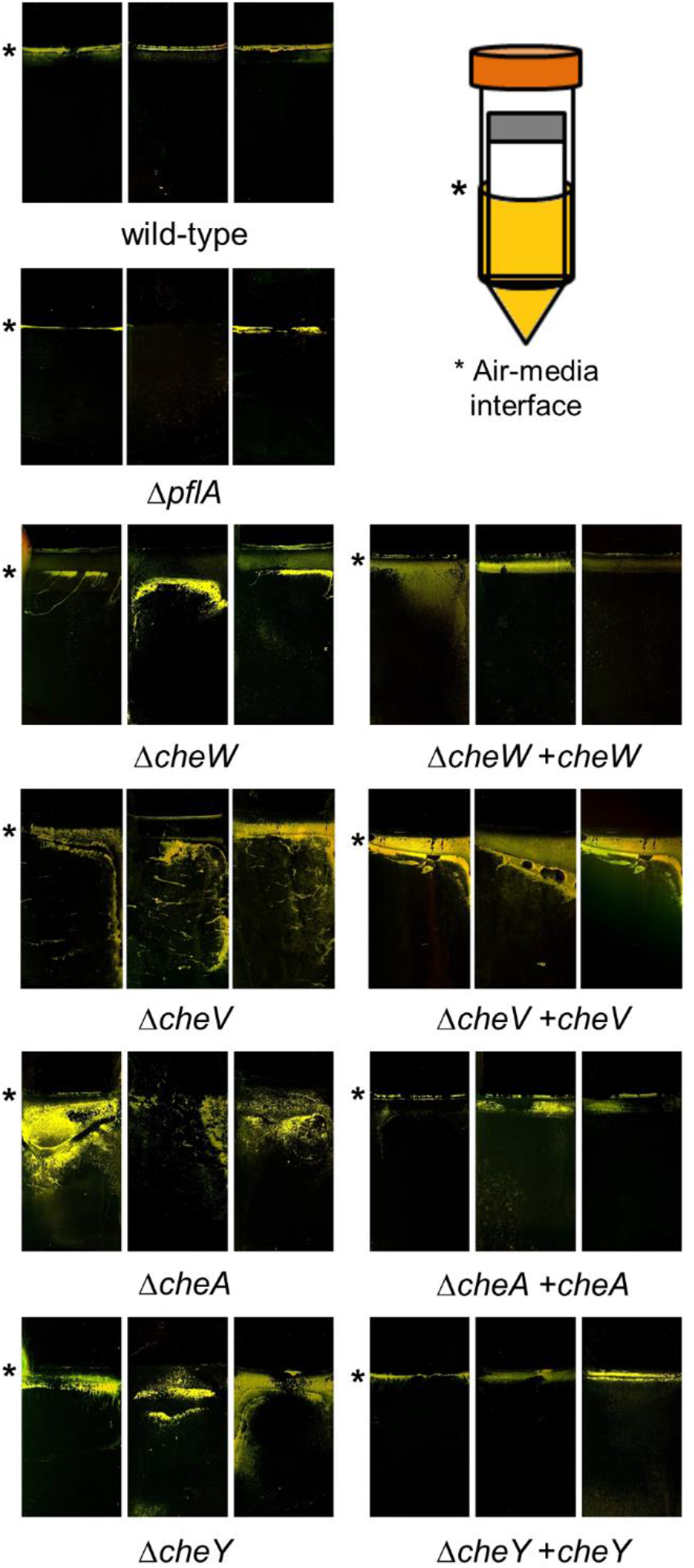
Chemotaxis is required for biofilm formation at the air-media interface. Static cultures of wild-type strain, chemotaxis mutants and complemented mutants were grown in the presence of a sterile glass slide, as shown in the cartoon on the top right (37°C, microaerobic, 48 hrs). As a control, the paralyzed flagella strain (Δ*pflA*) was included, which produces flagella but is non-motile. Glass slides were removed and stained with 1% crystal violet (CV). CV-stained biofilm was detected using a GenePix microarray scanner employing both 635 and 532 nm lasers. An asterisk shows the position of the air-media interface. Three independent biological repeats are shown for each strain.

## DISCUSSION

Flagellar motility plays an important role in colonisation and virulence of many bacterial pathogens. While the ability to move to or from locations is an important contributor to the infection process, such movement needs to be directed, and is commonly based on detection of external (e.g. chemical gradient) and internal stimuli (e.g. metabolic state). These stimuli can be positive or negative, with examples of positive stimuli being attractants such as nutrients, while negative stimuli can be repellents such as bile acids. The chemotaxis system integrates the signals from external and internal sensors through a signal transduction cascade consisting of MCPs, CheW/CheV, CheA and CheY, while other factors such as CheB or CheR, may modulate and fine-tune the signal transduction cascade, with diverse combinations observed throughout the bacterial kingdom (Lertsethtakarn *et al*, 2011; Micali and Endres, 2016). In this study we have dissected the core chemotaxis signalling pathway of the important foodborne zoonotic pathogen *C. jejuni*, which is the most common bacterial cause of gastroenteritis in the developed world. Directed motility is central to the process of intestinal colonisation by *C. jejuni*, as mutants defective in motility or chemotaxis are either unable to colonise or show significantly reduced efficiency of colonisation (de Vries *et al*, 2017; Yao *et al*, 1997).

While the CheA and CheY proteins are universal to bacterial chemotaxis systems, and bacteria may have one or multiple copies of these proteins, there is considerable variation in the adapter proteins such as CheV and CheW. *Campylobacter* is not exceptional in having both CheV and CheW proteins. Although CheW is the most common chemotaxis ‘adaptor’ protein (found in nearly 100% of genomes containing the core chemotaxis components, CheV was found in almost 40% of chemotaxis-positive genomes (Wuichet and Zhulin, 2010). Only three genomes (*B. thuringiensis*, *B. weihenstephanensis*, and *L. monocytogenes*) were found to contain CheV orthologs without CheW orthologs, suggesting that CheV alone can function in the MCP-CheA complex (Wuichet and Zhulin, 2010). In *C. jejuni*, the chemotaxis phenotype of the Δ*cheW* mutant is the least severe (Figures 2, 3 and 5), suggesting that CheV does function to elicit some signal transduction event.

Similar effects were observed in *B. subtilis* (Rosario *et al*, 1994) and further showed that the CheW domain of CheV was sufficient for signal transduction. A direct protein-protein interaction between CheV and Tlp4 (deoxycholate sensor, Cj0262c), Tlp6 (TCA intermediates sensor, Cj0448c), and Tlp8 (redox sensor CetZ) has been documented (Parrish *et al*, 2007) and Yeast Two- and Three-Hybrid analysis and immunoprecipitation were used to show that Tlp1 interacts with both CheW and CheV, although the interaction with CheV was much stronger and localised to the CheV-W domain (Hartley-Tassell *et al*, 2010). Both Δ*cheV* and Δ*cheW* mutants had chemotactic motility defects in Brucella media although chemotaxis towards aspartate was still competent in either mutant (Hartley-Tassell *et al*, 2010). Similarly, the novel chemotaxis protein Cj0371 of *C. jejuni* was shown to interact with the CheV protein, and influence ATPase activity of CheA (Du *et al*, 2016), suggesting that multiple systems engage in the chemotaxis pathway to modulate signal transduction. An evolutionary genomics study showed that the presence of CheV is correlated with increased numbers of MCPs and proposed that CheV functions to accommodate signal transduction from a specific group of MCPs, exemplified by the *Salmonella* McpC chemoreceptor (Ortega and Zhulin, 2016). None of the ten *C. jejuni* MCPs share the architecture of the McpC-type sensor.

Moreover, *H. pylori* genomes encode only four chemotaxis sensors and three CheV proteins (Jimenez-Pearson *et al*, 2005). The C-terminal receiver domain on CheV most likely refines the function of this protein. It has previously been proposed that CheV acts as a phosphate sink, normalizing over-stimulation of CheA (Karatan *et al*, 2001; Ortega and Zhulin, 2016; Pittman *et al*, 2001).

Previous studies have implicated chemotaxis sensors in biofilm formation. A mutant lacking Tlp3 (Cj1564) showed increased biofilm formation (Rahman *et al*, 2014), while *cetZ* (Tlp8) mutants showed decreased biofilm formation (Chandrashekhar *et al*, 2015). We show that removing each component of the chemotaxis pathway results in disruption of biofilm formation at a discrete air-surface interface, and that this phenotype can be rescued by complementation for most of the chemotaxis genes investigated here. In this study, biofilm formation was reduced at the air-media interface in the Δ*cheV* and Δ*cheW* mutants, while *cheV and cheW* mutants of *C. jejuni* have also been reported to have increased biofilm formation (Tram *et al*, 2020). While this may appear conflicting, there are considerable differences in experimental setup. The well-based biofilm assay may also measure agglutinated cells stuck to the bottom of the well, whereas the air-media interface assay used in our study only qualitatively investigates the cells at that interface, and does not include cells congegrating at the bottom of wells. Many assays used for measuring biofilm formation are relatively crude, need careful interpretation to avoid over-interpretation, and are difficult to directly compare. However, these minor differences do not alter the conclusion that the chemotaxis system likely has a role in coordinating biofilm formation in *C. jejuni*, and that this coordinating role may extend to shedding and dispersal of cells from the biofilm. This study is therefore the first to present a role for signal transduction in the active dispersal of cells from a *C. jejuni* biofilm. In food-processing environments, biofilms present a reservoir of cells that can subsequently re-contaminate the food chain highlighting the need to learn more about biofilm dispersal.

In conclusion, the ability to couple flagella rotation with environmental sensing is an effective adaptive mechanism allowing bacteria to seek an optimum environment. While chemotaxis systems from model organisms such as *E. coli* and *B. subtilis* provide an effective model for studying signal transduction in non-paradigm organisms, species-specific modulations and augmentations abound and require focussed investigation. Such genus- and species-specific analyses will be required to better understand chemotaxis in other Epsilon-proteobacteria and other organisms that share elements of the *Campylobacter* chemotaxis system and provides a further paradigm for chemotaxis signal transduction.

## Acknowledgments

The authors wish to thank Dr Henri Tapp for assistance with statistical evaluations, and members of the former IFR *Campylobacter* research group for helpful discussions. We would also like to thank Rachael Stanley for electron microscopy and Maddy Houchen and Robert Hindmarsh for microbiology media support.

