## Supplementary Information for "Inactivation of the core *cheVAWY* chemotaxis genes disrupts chemotactic motility and organised biofilm formation in *Campylobacter jejuni*" (from another version of this preprint: version 3, 2020-08-23)

**CONTENTS:**

Page 2: Figure S1

Page 3: Figure S2

Page 4: Table S1

Page 7: Table S2

Page 8: DOI of Supplementary Movie 1

Page 9: References

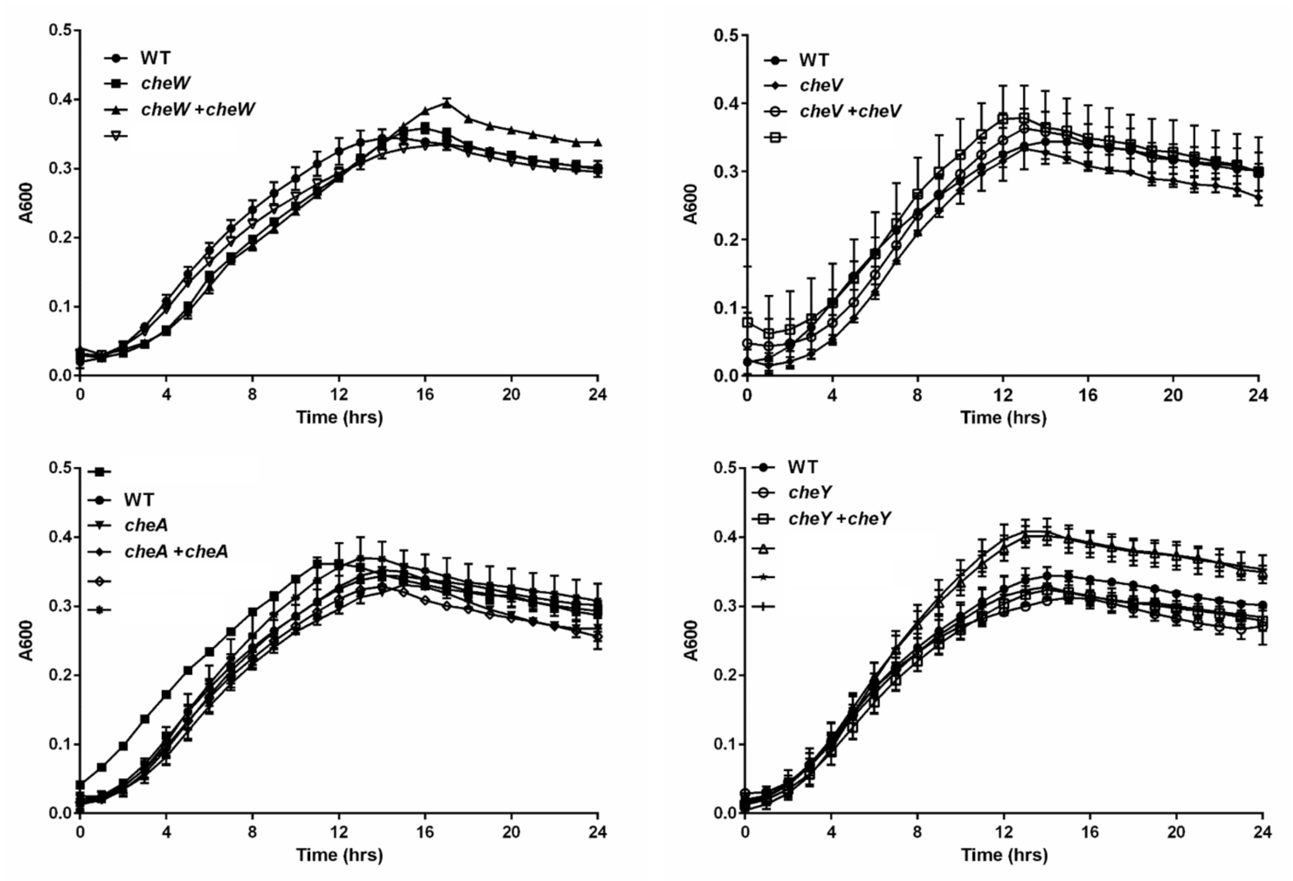


**Figure S1**. Chemotaxis mutants do not show impaired growth in Brucella media in microaerobic conditions. The graphs include additional mutants not used in this study.

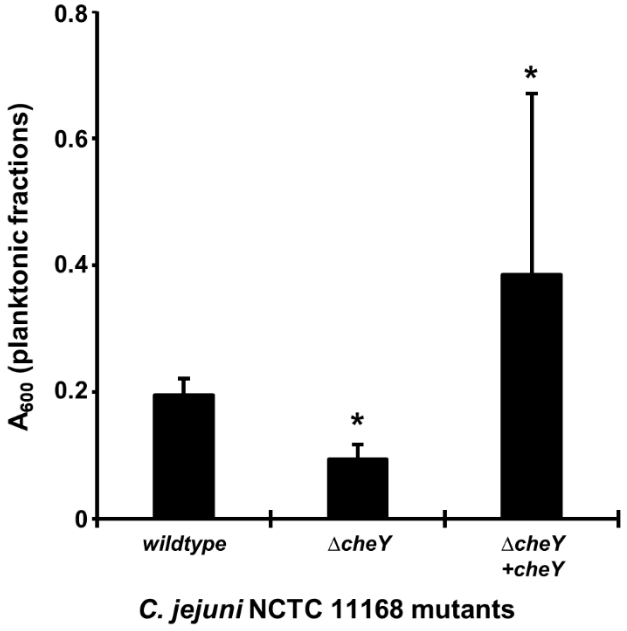


**Figure S2.** The biofilm phenotype of the Δ*cheY* mutant is linked to shedding of cells from the biofilm and the ability to swim to the air-media interface. To assess cell shedding from the biofilm, the $A_{600}$ of each planktonic culture used to prepare slide biofilm assays was measured. Planktonic cultures of at least three biological replicates were measured and error bars show standard deviation. An asterisk denotes statistically significant results, based on a Mann-Whitney test ($p<0.05$).

### Table S1. Oligonucleotide primers used in this study

| Name | Sequence [a] | Description |
|---|---|---|
| CheA-flankFwd | 5'-GCTTTAGATAAAGGTAccATTACAGG-3' | Upstream primer to amplify *cj0284c* plus ≈500bp upstream flanking sequence. Introduces *Kpn*I site. |
| CheA-flankRev | 5'-GATGGTTTCAgCATgCATTAAAGTAGC-3' | Downstream primer to amplify *cj0284c* plus ≈500bp downstream flanking sequence. Introduces *Sph*I site. |
| cheY_flankFwd | 5'-CCAAACggTaCCATAATAATACCC-3' | Upstream primer to amplify *cj1118c* plus ≈500bp upstream flanking sequence. Introduces *Kpn*I site. |
| cheY_flankRev | 5'-CGTTTGGTATAcTgcaGCTTCGC-3' | Downstream primer to amplify *cj1118c* plus ≈500bp downstream flanking sequence. Introduces *Pst*I site. |
| cheW_flankFwd | 5'-GGGTGATTgaaTtCAAAATATCC-3' | Upstream primer to amplify *cj0283c* plus ≈500bp upstream flanking sequence. Introduces *Eco*RI site. |
| cheW_flankRev | 5'-GCACAAAcTGCAgTCTTTAATCC-3' | Downstream primer to amplify *cj0283c* plus ≈500bp downstream flanking sequence. Introduces *Pst*I site. |
| cheV_flankFwd | 5'-CCTTACATgGtACcTTTAGAAGAATTGG-3' | Upstream primer to amplify *cj0285c* plus ≈500bp upstream flanking sequence. Introduces *Kpn*I site. |
| cheV_flankRev | 5'- GCTGTATTGTTAGcatGcTTAGGTGC -3' | Downstream primer to amplify *cj0285c* plus ≈500bp downstream flanking sequence. Introduces *Sph*I site. |
| cheAinverseA | 5'-CTAGGGATCCGCTCTACTAAATCATGGTCG-3' | N-terminal inverse PCR primer for replacing *cj0284c* with an antibiotic cassette. |
| cheAinverseB | 5'-CTAGGGATCCCCAGAATACTTGGAAAATGTGG-3' | C-terminal inverse PCR primer for replacing *cj0284c* with an antibiotic cassette. |
| cheYinverseStart | 5'-CTAGGGATCCCGTGTCCAAGTCTTGTTAGG-3' | N-terminal inverse PCR primer for replacing *cj1118c* with an antibiotic cassette. |
| cheYinverseEnd | 5'-CTAGGGATCCGGAAGTGGAGAAGGTGCAGC-3' | C-terminal inverse PCR primer for replacing *cj1118c* with an antibiotic cassette. |
| cheWinverseStart | 5'-CTAGGGATCCGCTGTTTTTGCAAAATTTGC-3' | N-terminal inverse PCR primer for replacing *cj0283c* with an antibiotic cassette. |
| cheWinverseEnd | 5'-CTAGGGATCCGTAGAAGCTTTGCTTAAGCG-3' | C-terminal inverse PCR primer for replacing *cj0283c* with an antibiotic cassette. |
| cheV_inverseA | 5'-GCATGGATCCGCTCCATTTCATTTGAACCC-3' | N-terminal inverse PCR primer for replacing *cj0285c* with an antibiotic cassette. |
| cheV_inverseB | 5'-GCATGGATCCCGATTTCTTCAATGAAATTGC-3' | C-terminal inverse PCR primer for replacing *cj0285c* with an antibiotic cassette. |
| cheAcompA | 5'-GGTAAAccATGGAAGATATGCAAG-3' | Primer used to amplify *cj0284c* for complementation in pKfdxA |
| cheAcompB | 5'-GCTCTAATTTTcCATggCTCATATTC-3' | Reverse primer used to amplify *cj0284c* for complementation in pKfdxA. |
| cheYcompA | 5'-GGACGAAATcaTGAAATTG-3' | Primer used to amplify *cj1118c* for complementation in pCfdxA. The start codon is changed from valine (native sequence) to methionine. |
| cheYcompB | 5'-GCATTTAGAATcaTgAAAATCAG-3' | Reverse primer used to amplify *cj1118c* for complementation in pCfdxA. |

| | | |
|---|---|---|
| cheWcompA | 5'-GGATAAGAtcATGAGTAATGAAAAATTAG-3' | Primer used to amplify *cj0283c* for complementation in pCfdxA |
| cheWcompB | 5'-GCATCAAAGTCAtgAACACAAAG-3' | Reverse primer used to amplify *cj0283c* for complementation in pCfdxA. |
| cheV_compFwd | 5’-CAGAAATTGAGGGGTCAAcATGTTTGATG-3’ | Primer used to amplify *cj0285c* for complementation in pCfdxA |
| cheV_compRev | 5’-GCATATCTTaCATgTTTTACCCCTGTTCTTG-3’ | Reverse primer used to amplify *cj0285c* for complementation in pCfdxA. |
| 0046Fcheck3 | 5’-GCAGAGCACTTGATTTTAGTGTGTGC-3’ | Primer for checking inserts into pseudogene *cj0046* used for complementation (upstream primer). |
| 0046Rcheck | 5’-CCTGGAGAAGTATTAGATAGTAGCGG-3’ | Primer for checking inserts into pseudogene *cj0046* used for complementation (downstream primer). |
| KmReadOut | 5'-CGGGGAAGAACAGTATGTCGAGC-3’ | Reads out from the 3’ end of the kanamycin resistance cassette. |
| KmPrReadOut | 5’-GCGATATCTTCTATATAAGCGTACCG-3’ | Reads out from the 5’ end of the kanamycin resistance cassette. |
| CatReadOut | 5’- CGTTTGTGACGGCTTTCATGTTTGC-3’ | Reads out from the 3’ end of the chloramphenicol resistance cassette. |
| CatPrReadOut | 5’-GGTCGAAATACTCTTTTCGTGTCC-3’ | Reads out from the 5’ end of the chloramphenicol resistance cassette. |
| 0046FwdSeq | 5’-GCTATGTTTATTACAGCTACAGC-3’ | Primer for sequencing inserts in pCfdxA/pKfdxA. |

[a] Restriction enzyme sites that are introduced for cloning purposes are underlined. Mismatched bases are in lowercase.

**Table S2.** Plasmids used in this study.

| **Plasmids** | **Description** [a] |
|---|---|
| pNEB193 | General cloning vector (NEB) |
| pCfdxA | Complementation plasmid containing *cj0046* flanks, chloramphenicol resistance cassette and *fdxA* promoter (Reuter and van Vliet, 2013) |
| pKfdxA | Complementation plasmid containing *cj0046* flanks, kanamycin resistance cassette and *fdxA* promoter (Reuter and van Vliet, 2013) |
| pCASO27F | *cheY* (*cj1118c*) disruption plasmid, $kan^R$ |
| pCASO28F | *cheW* (*cj0283c*) disruption plasmid, $kan^R$ |
| pCASO54R | *cheA* (*cj0284c*) disruption plasmid, $cat^R$ |
| pCASO69F | *cheV* (*cj0285c*) disruption plasmid, $kan^R$ |
| pCASO59F | *cheY* complementation plasmid, $cat^R$ |
| pCASO60F | *cheW* complementation plasmid, $cat^R$ |
| pCASO58F | *cheA* complementation plasmid, $kan^R$ |
| pCASO72F | *cheV* complementation plasmid, $cat^R$ |
| pMARKan9 | Source of kanamycin resistance cassette (Reuter *et al*, 2015) |
| pTopCat | Source of chloramphenicol resistance cassette (van Vliet *et al*, 1998) |

a. $kan^R$, kanamycin resistance; $cat^R$, chloramphenicol resistance

### SUPPLEMENTARY MOVIES

**Supplementary Movie 1.** Swimming behaviour of the *C. jejuni cheY* mutant.

Available via FigShare, doi: 10.15126/surreydata.7530881
